# Brain Antibody Sequence Evaluation (BASE): an easy-to-use software for complete data analysis in single cell immunoglobulin cloning

**DOI:** 10.1101/836999

**Authors:** S. Momsen Reincke, Harald Prüss, Jakob Kreye

## Abstract

**Background:** Repertoire analysis of patient-derived recombinant monoclonal antibodies is an important tool to study the role of B cells in autoimmune diseases of the human brain and beyond. Current protocols for generation of patient-derived recombinant monoclonal antibody libraries are time-consuming and contain repetitive steps, some of which can be assisted with the help of software automation.

**Results:** We developed BASE, an easy-to-use software for complete data analysis in single cell immunoglobulin cloning. BASE consists of two modules: aBASE for immunological annotations and cloning primer lookup, and cBASE for plasmid sequence identity confirmation before expression. Comparing automated BASE analysis with manual analysis we confirmed the validity of BASE output: identity between manual and automated aBASE analysis was 100% for all outputs, except for immunoglobulin isotype determination. In this case, aBASE yielded correct results in 96% of cases, whereas 4% of cases required manual confirmation. cBASE automatically concluded expression recommendations in 89.8% of cases, 91.8% of which were identical to manually derived results and none of them were false-positive.

**Conclusions:** BASE offers an easy-to-use software solution suitable for complete Ig sequence data analysis and tracking during recombinant mcAB cloning from single cells. Plasmid sequence identity confirmation by cBASE offers functionality not provided by existing software solutions in the field and will help to reduce time-consuming steps of the monoclonal antibody generation workflow.

## Background

Repertoire analysis of patient-derived recombinant monoclonal antibodies (mcAB) has become a state-of-the-art approach to investigate B cell (patho-)physiology and assess the role of (auto-)antibodies in disease progression in a variety of autoimmune and infectious diseases ([1],[2],[3]). Full repertoire data provide information about genetic composition for each single cell immunoglobulin (Ig), allowing identification of expanded Ig clones and providing insights into restrictions or enrichments of Ig gene usage in B cell populations ([4]). Moreover, recombinant expression approaches can generate libraries of purified mcABs which can be directly used in functional assays and further downstream applications.

In neuroscience, applying these approaches to cerebrospinal fluid (CSF)-derived cells has opened a rapidly developing field: Studying central nervous system mcAB repertoires (brain antibody-omics) enables insights into the physiological or pathogenic role of (auto-)antibodies in health and disease. First CSF cell studies showed direct pathogenic effects of single autoimmune encephalitis patient-derived mcABs in vitro and in vivo ([5], [6], [7], [8]), and also indicate local enrichment of disease-specific antibody-producing cells. Besides studying disease mechanisms, each CSF-derived mcAB can potentially be used for the development of novel diagnostic or therapeutic applications ([9], [10]), strongly promoting further investigations of mcAB repertoires in a broader spectrum of neuropsychiatric diseases. As CSF cell samples are challenging to obtain and typically have lower cell counts than samples from other body compartments, recombinant CSF cell cloning is naturally limited. Thus, protocols require optimization to increase successful expression rates from derived single cells in comparison to bulk approaches from blood samples.

Existing solutions for antibody sequence data analysis allow high-throughput processing in a database environment and are typically designed to work with next generation sequencing (NGS) ([11], [12], [13],[14]). These tools are designed to perform clonality analysis of bulk datasets. With the exception of sciReptor ([14]), they are not designed to handle single-cell data, therefore Ig heavy and light chain pairing is not preserved, and expression of all cloned antibodies is not possible.

In CSF mcAB studies, clonality analysis is less meaningful due to the low number of available cells compared to blood. We and others therefore focus on the identification and characterization of single mcABs with reactivity against a given auto-antigen. To this end, we identified three requirements for a software-assisted automated cloning pipeline, which are not yet fulfilled by available software solutions: (I) easy-to-access data in a human readable format; (II) data processing and cloning recommendations for all analyzed sequences in order to facilitate and optimize cloning efficiency from limited cell numbers, and (III) automated sequence data comparison from cloned plasmid genes to cDNA-derived sequences.

To address these requirements, we developed BASE (Brain Antibody Sequence Evaluation), an easy-to-use software solution suitable for complete Ig sequence data analysis and tracking during recombinant mcAB cloning from single cells.

## Implementation

### Design priorities and user workflow

BASE is designed to be easy to use, work with human readable formats whenever possible, be cross-platform, have minimal dependencies and make use of a modular design. It is designed around a core library (libBASE). The user interface is provided by two scripts, corresponding to the two relevant use cases in monoclonal antibody generation (Fig. 1). From amplified single cell cDNA, Ig sequences are obtained through Sanger sequencing, from which primary analysis and recommendations are provided for specific gene cloning using the aBASE (analyze-BASE) interface. Ig variable parts are then cloned into Ig expression vectors using a PCR-based approach. This cloning step can introduce nucleotide differences into the expression plasmid in comparison to the amplified single cell cDNA. Before recombinant expression of the mcAB, cBASE (compare-BASE) therefore confirms the identity of expression vector plasmid sequence and amplified cDNA, followed by the evaluation of suitability for expression. Sequence analysis algorithms for cBASE were designed to have a high positive predictive value for expression recommendation, i.e. to only recommend expression of a plasmid if it is 100% identical with the corresponding amplified cDNA.

**Figure 1:**
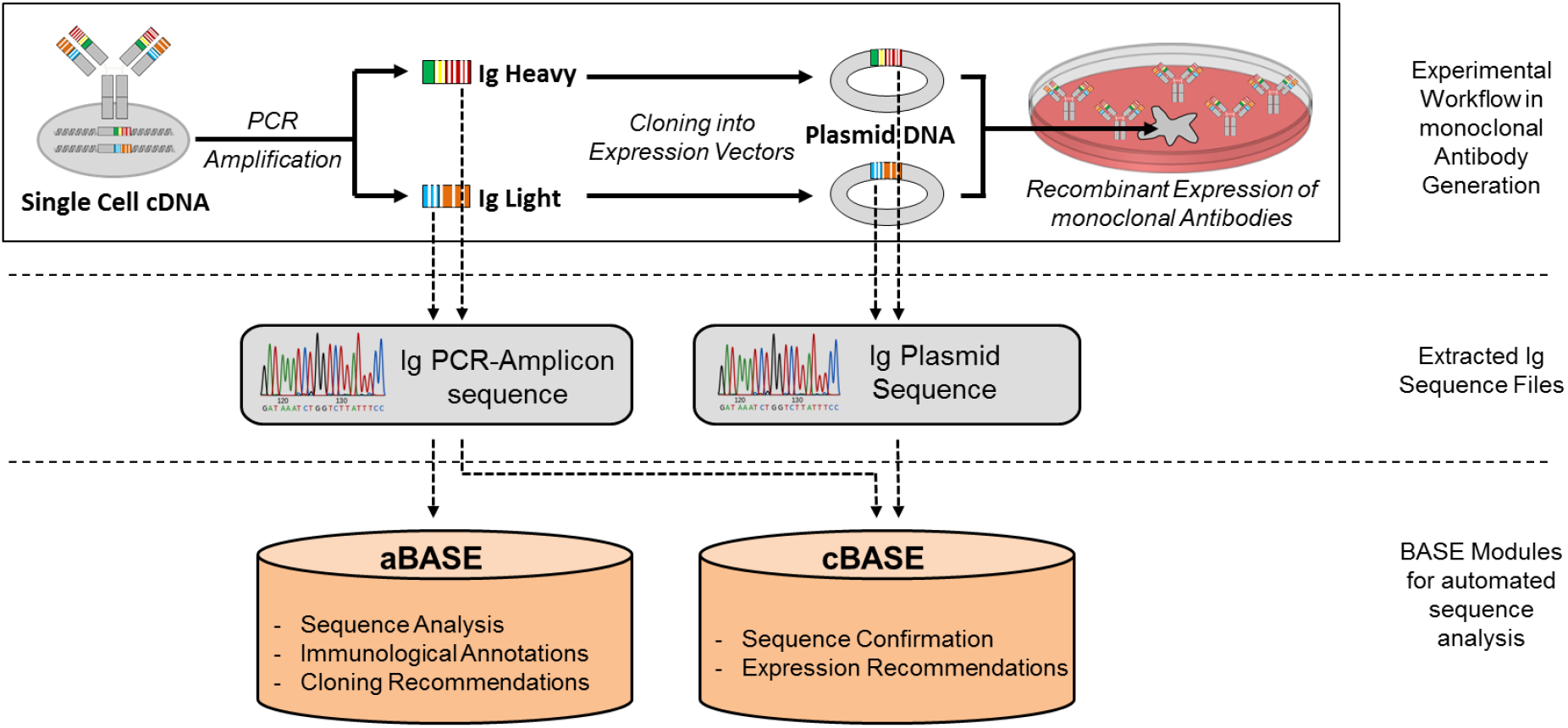
Monoclonal antibody generation workflow with BASE assistance. **A:** Overview of the experimental procedure (top row), corresponding raw sequencing files (middle) and BASE modules (bottom). Ig = immunoglobulin.

### Description of the core module libBASE

Single-cell Sanger sequencing data are loaded on a per-file basis and checked for sufficient quality for downstream analysis by excluding sequencing reads with Phread quality lower than 12 and sequencing read smaller than 50 nucleotides. IgBLAST ([15]) is used for alignment of single-cell sequence data with human Ig germline database sequences. Best matching immunoglobulin V, D, and J genes used as well as gene functionality, somatic hypermutation (SHM) count and full amino acid sequence and length of complementarity-determining region 3 (CDR3) are extracted. Furthermore, the Ig isotype is determined using BLAST ([16] to identify the best matching human constant Ig sequence ([14]). If this is inconclusive, typically due to either short sequence reads or low sequencing quality, the Ig isotype is determined using a heuristic customized to our primers. Suitable cloning primers for subsequent expression cloning are identified in a customizable table.

### Description of the aBASE user interface

All relevant data acquired during mcAB generation is stored in an .xlsx table, in which each line represents one single cell with a unique mcAB identifier. This format allows the connection of sequence data derived information with any single cell metadata from other sources (e.g. cell population assignment from FACS analysis), which can simply be added in any customized extent and order. aBASE takes this .xlsx file as an input and initiates a standardized analysis algorithm for each sequence file using the core module libBASE. An output file is then automatically generated which combines the input information with the acquired data complemented by individual recommendations for specific gene cloning such as the suitable gene-specific cloning primers and the presence of additional restriction sites (Fig. 2). Moreover, aBASE feedbacks the reliability of its analysis results in a comment column, suggesting in some cases a manual re-inspection of the data or re-sequencing of the sample, typically related to low sequencing quality.

**Figure 2:**
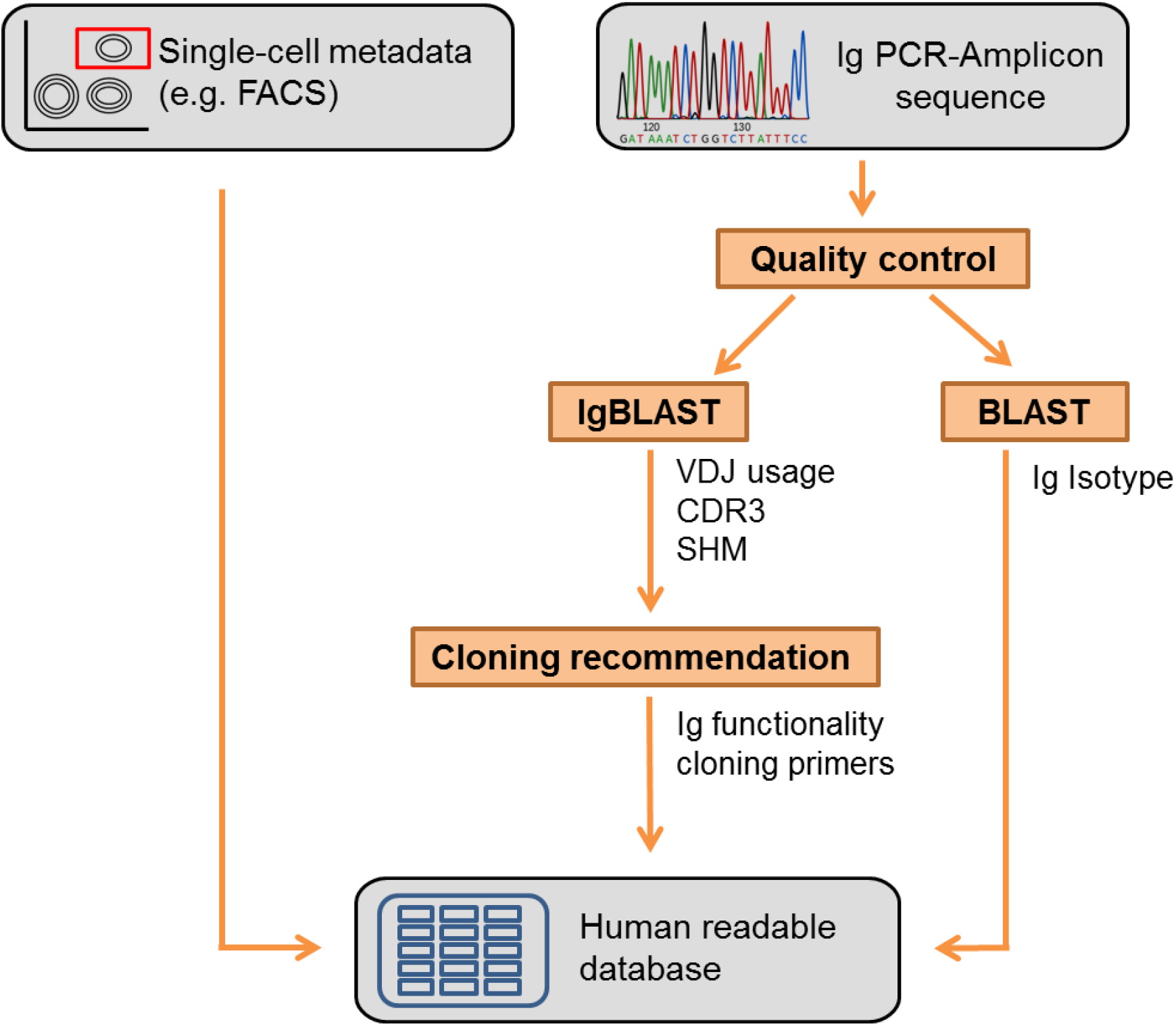
Workflow and implementation of aBASE: After quality control, PCR-amplicon sequencing files are parsed, blasted against germline VDJ-genes using IgBLAST, and blasted against constant gene parts using BLAST. Output is used to generate cloning recommendations and written to a human-readable database. FACS = fluorescence activated cell sorting. CDR = complementarity determining region. SHM = somatic hypermutations. Ig = immunoglobulin.

### Description of the cBASE user interface

After Ig genes have been cloned into the respective expression vectors containing the constant Ig domain, plasmids are sequenced using a primer annealing upstream of the start codon. cBASE aligns and compares the plasmid Ig sequence with the amplified cDNA-derived Ig sequence by displaying nucleotide differences, assigns their relevance for amino acid changes and evaluates possible influences of primer binding interactions. Only plasmids with 100% amino acid sequence identity with the respective amplified cDNA are supposed to be released for expression. Results of this analysis are presented in a color encoded format, allowing rapid differentiation of plasmid genes suitable for mcAB expression (green), clones of which further bacterial colonies need to be analyzed for suitability (red) and genes where manual sequence inspection is recommended (brown). The latter typically occurs when sequence quality is low and automated nucleotide assignment may contain errors (Fig. 3). Additionally, the total SHM count in the variable immunoglobulin V-gene is separately displayed for the amplified cDNA and plasmid Ig sequence read, allowing an easy correction of aBASE-derived SHM numbers as some may falsely be accounted from primer binding interactions in the beginning of the amplified cDNA sequence read.

**Figure 3:**
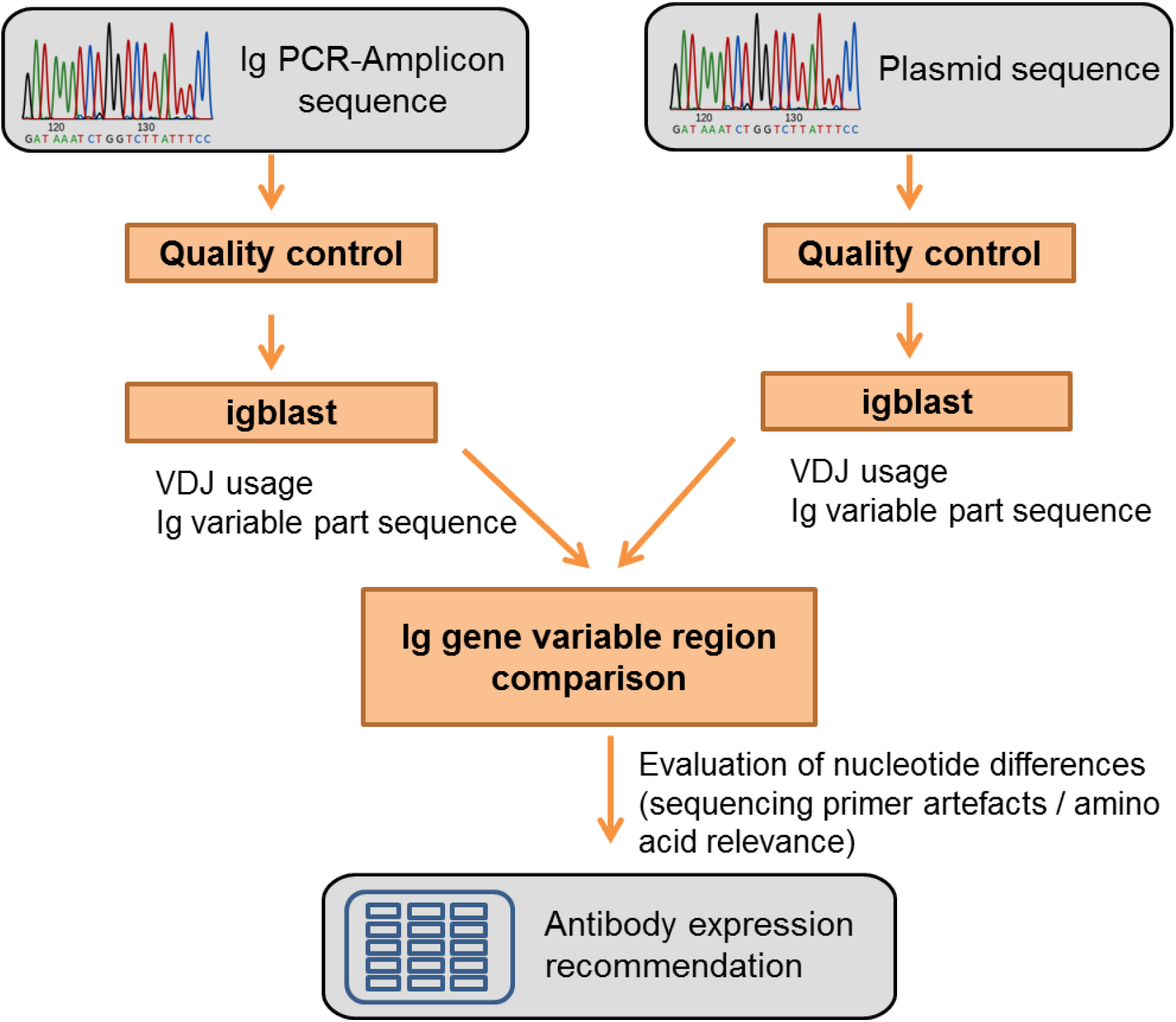
Workflow and implementation of cBASE: After quality control, PCR-amplicon and plasmid sequencing files are parsed, and then blasted against germline VDJ-genes using IgBLAST. Ig gene variable regions are compared nucleotide by nucleotide and antibody expression is recommended if no amino acid changing mutations are detected. CDR = complementarity determining region. Ig = immunoglobulin.

### Code dependencies and availability

BASE is written in python and requires local installation of IgBLAST and BLAST. The source code is published under GNU General Public License (GPL-v3) and available under https://github.com/automatedSequencing/BASE.

## Results

To evaluate the validity of our automated analysis using the two BASE software interfaces, we performed automated BASE analysis from a recently acquired mcAB sequence data set, which previously had been analyzed manually, and compared the results. As differences could potentially be attributed to software bugs or errors in previous manual inspection, divergent data were re-inspected manually. The data set is derived from a CSF cell sample processed using mcAB repertoire cloning in our laboratory (sample ID #AI ENC 113, Kreye et al. in preparation), including a total of 181 amplified cDNA Ig sequence reads for aBASE validation and a total of 176 plasmid Ig sequence reads for cBASE validation.

### Validation of primary Ig sequence evaluation using aBASE

Comparison of the results of the analysis performed by aBASE and the manual analysis expectedly showed no relevant differences in the parameters belonging to pre-processing analysis (quality value and length of the sequence read) and the data extracted from IgBLAST (best matching germline Ig genes, gene functionality, somatic hypermutation count, CDR3 sequence and length) (Table 1). The only differences resulted from manual typing inaccuracies and were not counted as automated sequencing analysis mistakes. Ig isotype identification is performed from Ig heavy chain sequences only and is commonly hindered by low quality at the initial part of the sequence read and annealing of multiple primers, therefore being the most delicate part of Ig gene analysis. aBASE correctly identified the Ig isotype in 92 out of 96 cases (95.8%, Table 1) without requiring any further user action. In 2 cases the automated analysis yielded the correct Ig isotype but flagged it as “low-confidence” and recommended manual sequence inspection. In the remaining 2 cases (2.1%), Ig isotype could not be determined automatically. Correct cloning recommendations were derived in all cases.

**Table 1:**
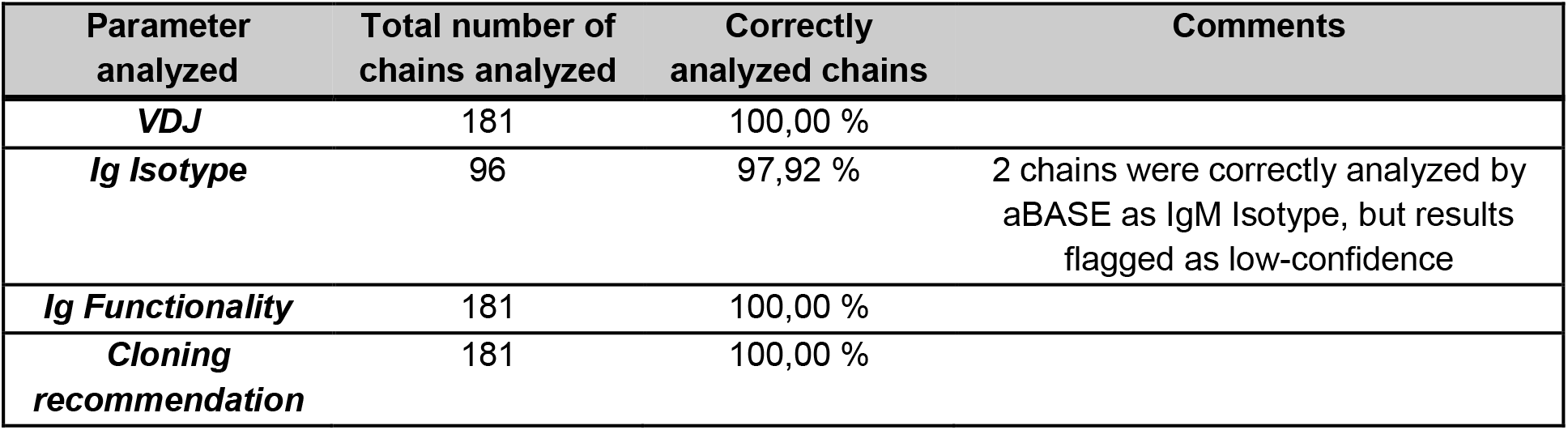
Validation of aBASE. Number of correctly identified parameters using automated aBASE evaluation compared to the results of manual analysis.

| Parameter analyzed | Total number of chains analyzed | Correctly analyzed chains | Comments |
| --- | --- | --- | --- |
| **VDJ** | 181 | 100,00 % |  |
| **Ig Isotype** | 96 | 97,92 % | 2 chains were correctly analyzed by aBASE as IgM Isotype, but results flagged as low-confidence |
| **Ig Functionality** | 181 | 100,00 % |  |
| **Cloning recommendation** | 181 | 100,00 % |  |

### Validation of plasmid versus amplified cDNA Ig sequence comparison using cBASE

Automated cBASE analysis concluded recommendations for expression suitability in 158 out of 176 cases without requiring user interaction (89.8%, Table 2). Manual re-evaluation was required in 18 cases (10.2%), associated with low sequence read quality (e.g. yielding putative frameshifts), which naturally limits automated analysis. In 145 out of 158 cases (91.8%; Table 2), cBASE algorithm derived conclusions were considered correct which occurred if cBASE either a) correctly concluded no relevant differences between plasmid and amplified cDNA or b) correctly identified relevant differences and dismissed the plasmid as not suitable for expression. Of the 13 cases in which automated analysis came to different conclusions than manual evaluation, there were no false-positives, i.e. cBASE never advised to use a plasmid for expression that did not fulfill requirements for expression suitability in manual inspection. Instead, in all 13 differing cases, the software evaluation was technically correct as there were relevant nucleotide differences in the sequencing results. However, those differences were likely not reflecting changes in the DNA sequence, but rather related to poor sequence quality in parts of the amplified cDNA Ig sequence. Nonetheless, manual assessment suggested functional genes which were chosen for expression, thus those cBASE recommendations were considered false-negative. Overall, BASE expression recommendations in this study had a positive predictive value of 100% and a negative predictive value of 76.4%.

**Table 2:** Validation of cBASE. 176 expression plasmids were compared with the corresponding amplified cDNA sequences. An expression recommendation by cBASE was considered false-negative if the software recommended against using the plasmid, although upon manual inspection the differences between plasmid and amplified were negligible (and most likely due to sequencing quality artifacts).

| **cBASE analysis results** | **# of chains** | **Comments** |
| --- | --- | --- |
| ***correct expression recommendation*** | 145/176 |  |
| ***false-negative expression recommendation*** | 13/176 | mostly related to punctual low sequencing quality either in the plasmid or in the amplified cDNA Ig sequence |
| ***no automatic recommendation*** | 18/176 | mostly related to general low sequencing quality either in the plasmid or in the amplified cDNA Ig sequence |
| ***false-positive recommendation*** | 0/176 |  |

## Discussion

With BASE we have established an easy-to-use software solution to facilitate mcAB sequence analysis and cloning in the process of monoclonal antibody generation. Automated sequence analysis using aBASE or plasmid expression recommendations using cBASE save time (about 3-5 minutes per sequence), have low error rates, and the requirement for user re-analysis is limited to cases with poor sequence quality.

We validated aBASE and cBASE in a dataset which previously had been analyzed manually. The comparison of the results was generally favorable. We identified only a few minor differences in comparison to independent manual evaluation, which upon re-inspection were caused by typing errors during the initial human evaluation, further highlighting the benefits of automated sequence analysis by reducing human error. Moreover, our comparison indicates that cBASE expression recommendations have a positive predictive value of 100%, however, are limited in negative prediction (NPV of 76.4%). Our detailed re-analysis revealed that this occurs mostly when cDNA amplicon Sanger sequence has low quality. Therefore, cBASE performance may be modulated by introducing Sanger sequence quality thresholds for mcABs before Ig gene cloning. Higher quality threshold reduces the overall antibody yield and therefore may not be feasible if source cells are limited, as in the case of CSF-derived mcAB cloning. If so, we suggest a manual re-inspection of Ig plasmid sequences whenever cBASE yields a negative expression recommendation and cDNA amplicon Sanger sequence is of low quality.

BASE is optimized towards requirements from recombinant mcAB cloning from CSF cells, but can likewise be used with human B cells including antibody-secreting cells from any source, therefore enabling a much wider use in antibody-omics. Single cell-based approaches are most useful whenever sample access is difficult and cell count is limited (e.g. tumor cells, biopsy material, any body fluid other than blood). The workflow has been established from isolated single cells in a 96 well format, but under the same set up can similarly process nearly unlimited amounts of data points in one file, from any source of B cell population in any plate format. Also, multiple plates or specimens can be tracked and analyzed in one file without any further changes. Moreover, the BASE platform could easily be adapted to perform non-human Ig sequence evaluation with few adaptations in linking to databases for germline Ig genes and primers.

Planned future developments include implementation of additional modules to automatically import and link further data from the mcAB generation process other than sequence information into the same host file, such as flow cytometry single cell sort data, ELISA and microscopy data.

## Conclusions

The role of brain-targeting autoantibodies in neurological and psychiatric autoimmune and degenerative diseases is increasingly recognized, generating the need for more automated approaches in recombinant mcAbs production. BASE already resolves a bottleneck in today’s mostly non-automated workflow of antibody generation, but the full potential of BASE will be unfolded with upcoming advancements in the high-throughput sequencing as part of mcAB generation pipelines.

## List of abbreviations

CDR: complementarity determining region
CSF: cerebrospinal fluid
Ig: Immunoglobulin
mcAB: monoclonal antibody
NGS: Next-generation sequencing
PCR: Polymerase chain reaction
SHM: Somatic hypermutation

## Availability and requirements

Project name: BASE. Project home page: https://github.com/automatedSequencing/BASE. Operating system(s): Linux and Windows. Programming language: Python. Other requirements: IgBLAST and BLAST. License: GNU General Public License (GPL-v3).

## Acknowledgments

The authors would like to acknowledge the preceding work of Jordan Willis, who wrote pyigblast (https://github.com/andrejbranch/pyigblast) on which a part of BASE is based.

## Availability of data and materials

See https://github.com/automatedSequencing/BASE.

## Authors’ contributions

SMR implemented BASE and drafted the manuscript. HP conceived the project and edited the manuscript. JK conceived the project, drafted the manuscript and provided feedback on BASE functionality. All authors read and approved the final manuscript.

## References

1. Wardemann, H., et al., Predominant autoantibody production by early human B cell precursors. Science, 2003. 301(5638): p. 1374–7.

2. Scheid, J.F., et al., Broad diversity of neutralizing antibodies isolated from memory B cells in HIV-infected individuals. Nature, 2009. 458(7238): p. 636–40.

3. Murugan, R., et al., Clonal selection drives protective memory B cell responses in controlled human malaria infection. Sci Immunol, 2018. 3(20).

4. Schickel, J.N., et al., Self-reactive VH4-34-expressing IgG B cells recognize commensal bacteria. J Exp Med, 2017. 214(7): p. 1991–2003.

5. Bennett, J.L., et al., Intrathecal pathogenic anti-aquaporin-4 antibodies in early neuromyelitis optica. Annals of Neurology, 2009. 66(5): p. 617–629.

6. Kreye, J., et al., Human cerebrospinal fluid monoclonal N-methyl-D-aspartate receptor autoantibodies are sufficient for encephalitis pathogenesis. Brain, 2016. 139(Pt 10): p. 2641–2652.

7. Malviya, M., et al., NMDAR encephalitis: passive transfer from man to mouse by a recombinant antibody. Ann Clin Transl Neurol, 2017. 4(11): p. 768–783.

8. Jurek, B., et al., Human gestational NMDAR autoantibodies impair neonatal murine brain function. Ann Neurol, 2019.

9. Ly, L.T., et al., Affinities of human NMDA receptor autoantibodies: implications for disease mechanisms and clinical diagnostics. J Neurol, 2018. 265(11): p. 2625–2632.

10. Tradtrantip, L., et al., Anti-Aquaporin-4 monoclonal antibody blocker therapy for neuromyelitis optica. 2012. 71(3): p. 314–322.

11. Moorhouse, M.J., et al., ImmunoGlobulin galaxy (IGGalaxy) for simple determination and quantitation of immunoglobulin heavy chain rearrangements from NGS. BMC Immunol, 2014. 15: p. 59.

12. Vander Heiden, J.A., et al., pRESTO: a toolkit for processing high-throughput sequencing raw reads of lymphocyte receptor repertoires. Bioinformatics, 2014. 30(13): p. 1930–2.

13. Gupta, N.T., et al., Change-O: a toolkit for analyzing large-scale B cell immunoglobulin repertoire sequencing data. Bioinformatics, 2015. 31(20): p. 3356–8.

14. Imkeller, K., et al., sciReptor: analysis of single-cell level immunoglobulin repertoires. BMC Bioinformatics, 2016. 17: p. 67.

15. Ye, J., et al., IgBLAST: an immunoglobulin variable domain sequence analysis tool. Nucleic Acids Res, 2013. 41(Web Server issue): p. W34–40.

16. Altschul, S.F., et al., Basic local alignment search tool. J Mol Biol, 1990. 215(3): p. 403–10.

